# How many characters are needed to reconstruct a phylogeny?

**DOI:** 10.1101/2025.09.26.678777

**Authors:** Alessio Capobianco

## Abstract

Despite increased recent attention towards Bayesian phylogenetics and its applications in understanding macroevolutionary processes, it remains unclear how many discrete characters are needed to accurately estimate tree topologies in a Bayesian framework. This could be particularly relevant for morphological datasets used in phylogenetics, as they usually consist of few dozens to few hundreds of characters—orders of magnitude smaller than most molecular datasets.

I designed a simulation study in the software RevBayes to explore how the number of sampled discrete characters affects accuracy and precision of Bayesian phylogenetic estimates, under various setups differing in number of taxa, average number of state changes per character (i.e., tree length), and number of states per character. Results indicate that between 100 and 500 variable characters are necessary to reach sufficient accuracy and precision of phylogenetic estimates for as low as 20 tips. All other parameters being equal, multistate characters produce slightly more accurate estimates than binary characters, and more labile characters produce more accurate estimates for trees above 50 tips. The results of this study highlight the continuous need for global research efforts geared towards the characterization and digitization of interspecific morphological diversity in both extant and extinct taxa.

## 1. Introduction

We arguably live in a golden era of phylogenetics, with the past two decades characterized by: a dramatic increase in the availability of molecular sequences, and in the number of phylogenetic studies (1,2); the development of complex probabilistic models of character and lineage evolution (3–7); Bayesian approaches allowing for the modular integration of different data types (e.g., molecules, morphology, stratigraphy, biogeography) to derive time-calibrated trees (8,9); an ever-growing toolkit of phylogenetic comparative methods that harness phylogenetic estimates to answer all kinds of macroevolutionary questions (10–13). Alongside this recent explosion of phylogenetics and of its applications, considerably less attention has been devoted towards how dataset size affects the accuracy and precision of phylogenetic estimates. Limitations derived from dataset size might be particularly relevant for morphological phylogenetics, as morphological datasets usually range between few dozens to few hundreds of characters—considerably less than the thousands-to-millions of characters seen in molecular datasets.

Between the 1990s and the early 2000s, several papers discussed dataset size—in terms of both number of taxa and number of characters—mostly in the context of a broader debate about the role of morphological data in phylogenetics after the beginning of the molecular phylogenetics revolution (14– 22). These studies have important limitations that constrain their practical application in empirical phylogenetic research, including: the lack of a statistical framework that allows for simulation of characters under the same process assumed when estimating the phylogeny; the lack of metrics measuring accuracy of phylogenetic estimates (focusing rather on resolution and support, which do not measure how close an estimate is to the true phylogeny); the use of simulated datasets with unrealistic data sizes compared to most empirical morphological studies (either too small or too large); the absence of clear recommendations on the number of characters needed to reconstruct a phylogeny.

Knowing how many characters are needed to reconstruct a phylogenetic tree above a certain threshold of accuracy and precision is fundamental for two reasons: 1) it sets a target dataset size that can be helpful for elaborating a research plan and guiding data collection; 2) it defines realistic expectations on how far from the true phylogeny a certain phylogenetic hypothesis might be, given the size of the dataset used to reconstruct it.

Here, I designed a simulation study to explore how the number of sampled discrete characters affects accuracy and precision of Bayesian phylogenetic estimates, under various setups differing in number of taxa, average number of state changes per character (i.e., tree length), and number of states per character. The advantage of using a Bayesian framework is that data can be simulated under the exact same model and parameter settings used for phylogenetic inference, ensuring that the phylogenetic estimate would converge to the true phylogeny with data size approaching infinity. Thus, all potential discrepancies between estimated and true phylogeny will be solely due to the size of the dataset and the intrinsic stochasticity associated with it.

## 2. Material and methods

The simulation study was performed in the software RevBayes v1.2.3 (23) on the PalMuc high-performance computing cluster (HPC) at LMU Munich. To simulate discrete character evolution on phylogenetic trees, unrooted phylogenies were first simulated for various numbers of tips (5, 10, 20, 50, 100, or 200) and under different expected tree lengths—that is, different expected number of state changes per character across the whole tree (1, 3, or 10). Total tree length rather than individual branch lengths was picked as a control parameter, because it provides information on the properties of simulated characters independently of how many tips are included in the tree. Characters evolving on a tree with length = 1 will generate a pattern of states that is mostly consistent to either synapomorphic traits (derived traits shared by all members of a clade, corresponding to a single character state change on an internal branch) or autapomorphic traits (derived traits that are unique to a single lineage, corresponding to a single character state change on a terminal branch). On the contrary, characters evolving on a tree with length = 10 will mostly display homoplastic state patterns, with the same trait evolving multiple times independently along the phylogeny.

For each phylogeny, the topology was sampled from a uniform unrooted topology distribution, and each branch length was sampled from an exponential distribution with mean equal to the expected tree length divided by the number of tree branches. For each combination of numbers of tips and expected tree lengths, 50 different phylogenies were simulated, for a total of 6 (different numbers of tips) x 3 (different tree lengths) x 50 (replicates) = 900 phylogenies.

Discrete character data were simulated under each phylogeny using the Mkv model of morphological evolution (24). The Mkv model assumes that all characters evolve at the same rate, all character states are equiprobable, and only characters that are observed to be variable are sampled (24). When simulating under the Mkv model, the average effective number of state changes per character across the whole tree is slightly higher than the tree length, as characters that would be observed as invariant are not sampled (24,25). However, this does not change the qualitative assessment made above on the properties of characters simulated under different tree lengths.

The number of variable discrete characters simulated for each phylogeny varied between 20, 50, 100, 500, 1000, and 5000 characters. These values cover the whole range of dataset sizes that can be seen in empirical morphological phylogenetic analyses. Simulations of character evolution were repeated for 2-state (binary), 3-state, and 4-state characters, to explore how the number of character states—and thus the amount of information per character—can affect phylogenetic estimates. Table 1 summarizes the different parameters and parameter values used to simulate datasets. A total of 900 (phylogenies) x 6 (different numbers of characters) x 3 (different state spaces) = 16,200 datasets were simulated.

**Table 1.** List of all relevant parameters and their values used to simulate discrete character datasets.

| Parameter | Parameter values |
| --- | --- |
| Number of taxa | 5, 10, 20, 50, 100, 200 |
| Number of characters | 20, 50, 100, 500, 1000, 5000 |
| Expected tree length | 1, 3, 10 |
| Number of character states | 2, 3, 4 |
| Simulation replicates | 1–50 |

For each simulated dataset, a Bayesian phylogenetic analysis was performed in RevBayes. To ensure that there was no model misspecification, all model specifications and priors on parameter values were consistent with how the datasets were generated. Topology and branch length priors were set as identical to the distributions generating the simulated trees. The Mkv model was set as the substitution model, with a transition matrix (or Q-matrix) of appropriate size (2×2 for the binary datasets, 3×3 for the 3-state datasets, 4×4 for the 4-state datasets) (26). This mimics an ideal scenario in which the models used for phylogenetic inference are an accurate reflection of how the sampled characters evolved. For each analysis, four independent MCMC (Markov chain Monte Carlo) simulations were run for 100,000 iterations, sampling trees and parameters every 10 iterations. The first 10% of each MCMC simulation was discarded as burn-in. Convergence of continuous parameters between independent replicates for each simulated data analysis was assessed using the R package convenience (27), with thresholds for minimum effective sample size set as 200 and for the Kolmogorov-Smirnov test value calculated at precision level = 0.0177 (27). The posterior distribution of trees for each analysis was summarized into a maximum *a posteriori* (MAP) tree (in some literature called the highest posterior frequency tree (28)).

Three metrics were used to assess the accuracy and precision of phylogenetic estimates compared to the true tree under which data were simulated:

1. Normalized Robinson-Foulds (RF) distance (29) between MAP tree and true tree. This metric measures the accuracy of the single most probable tree in the posterior distribution, often used to summarize phylogenetic results and for downstream phylogenetic comparative analyses. Values below a threshold of 0.25 indicate a sufficiently accurate phylogenetic estimate. This metric was calculated in R using the package TreeDist v2.9.2 (30).
2. Percentage of true clades that are strongly supported. Clades are considered to be strongly supported if their posterior probability (PP) is above 0.95. This metric measures a combination of accuracy and precision of the phylogenetic estimate. Values above a threshold of 50% are indication of sufficient accuracy/precision. Posterior probabilities of true clades were calculated with the cladeProbability() method in RevBayes v1.2.3 (23).
3. Mean posterior probability of clades in the MAP tree. This metric measures the precision of the phylogenetic estimate. Values above a threshold of 0.75 indicate a sufficiently precise phylogeny. This metric was calculated in R using the package treeio v1.22.0 (31).

Threshold values for these metrics are necessarily subjective, as there are no universal standards employed in the field of phylogenetics to define when a phylogenetic estimate is “good enough”. For more detailed explanations about the chosen metrics and their selected threshold values, see the supplementary material. Phylogenetic estimates under a given combination of number of taxa, number of characters, number of character states, and tree length were considered to be sufficiently accurate and precise if the mean of all three metrics across simulation replicates passed the threshold values.

## 3. Results

Overall, accuracy and precision of phylogenetic estimates always increase with more characters (Fig. 1, Supplementary Figs). Nonetheless, how the number of characters impacts the quality of a phylogenetic estimate depends on the combination of all the other factors considered in this study: number of taxa, tree length, and (to a lesser degree) number of states.

**Figure 1.**
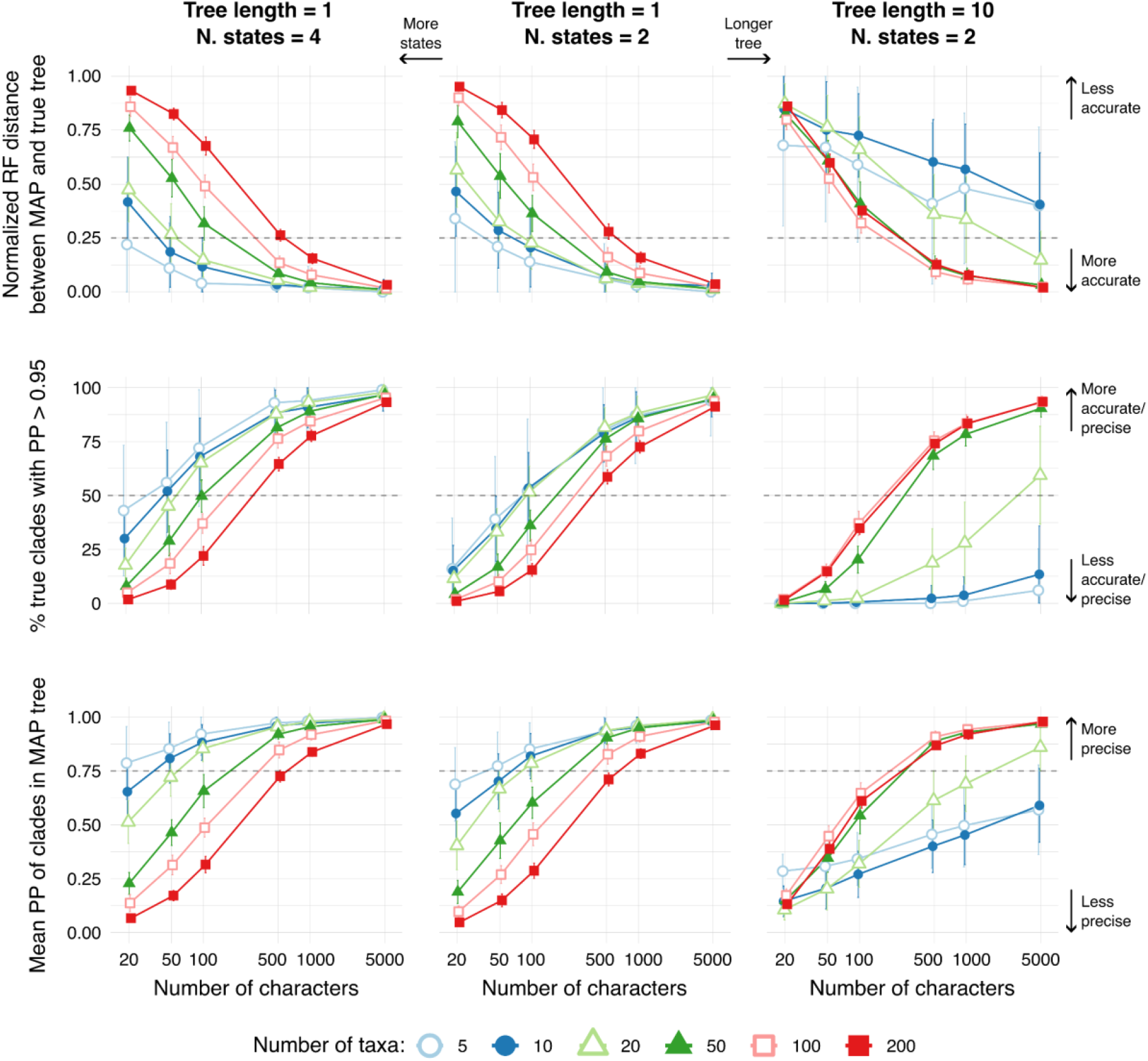
Accuracy and precision of Bayesian phylogenetic estimates for different combinations of number of characters, number of taxa, number of states per character, and total tree length. Metrics considered are: the normalized Robinson-Foulds (RF) distance between maximum *a posteriori* (MAP) tree and true tree (*top row*); the percentage of true clades that are strongly (PP > 0.95) supported (*middle row*); the mean posterior probability of clades in the MAP tree (*bottom row*). Horizontal dashed lines indicate threshold values for satisfactory levels of accuracy and precision. Mean (*symbol*) and standard deviation (*bars*) of the three metrics across simulation replicates are indicated. Tree length is expressed in units of expected state changes per character.

At tree length = 1 and keeping number of characters and states equal, trees with more taxa score worse than trees with fewer taxa in all three accuracy and precision metrics considered here (Fig. 1, Supplementary Figs). Thus, the larger the tree is in terms of number of taxa, the more characters are needed to reach satisfactory levels of accuracy and precision. Considering the mean of metrics across simulation replicates, threshold values are passed at 50 to 100 characters for up to 20 taxa, at 500 characters for 50–100 taxa, and at 1,000 characters for 200 taxa (Fig. 2). Increasing the number of states per character always has a positive effect, although this effect is almost negligible for larger trees.

**Figure 2.**
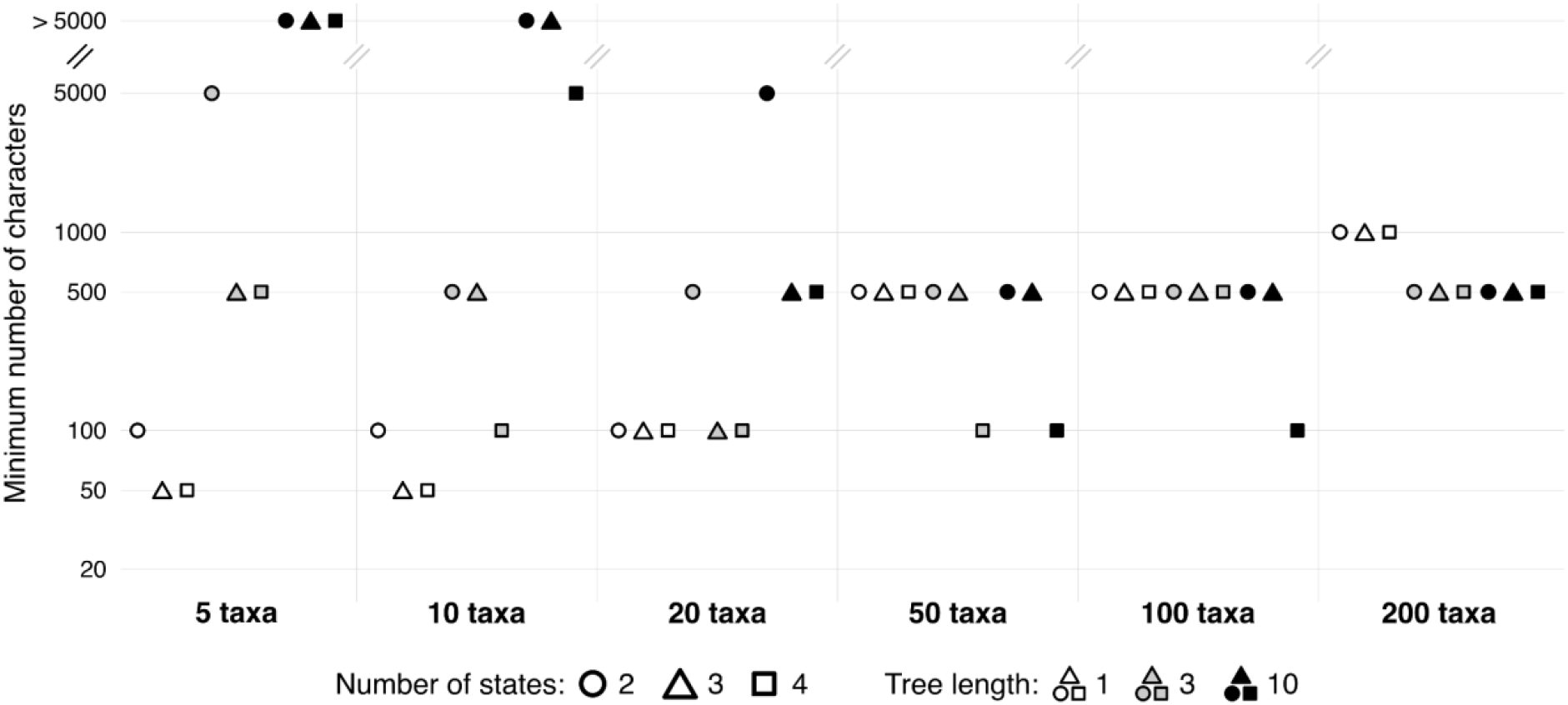
Minimum number of variable characters needed to reach satisfactory levels of accuracy and precision of the phylogenetic estimate, under different combinations of number of taxa, number of states per character, and total tree length. For each combination of those parameters, the mean across simulation replicates of all three metrics of accuracy and precision had to pass threshold values. Tree length is expressed in units of expected state changes per character.

At longer tree lengths, the negative correlation between number of taxa and accuracy/precision of the phylogenetic estimate breaks down into a more complex pattern. At tree length = 3, the normalized RF distance between MAP and true tree follows the same pattern seen for the shortest tree length only between 20 and 200 taxa, but not for less than 20 taxa (Supplementary Figs. 1–2). The accuracy of the phylogenetic estimate improves with an increased number of characters more slowly for 5–10 taxa than for 20 taxa or more. This means that at 500 characters or more, the normalized RF distance to the true tree for 5-taxa trees is similar or even higher than the one for 200-taxa trees. An analogous behaviour can be seen also for the other two metrics, with tree estimates for 5–10 taxa being less accurate and precise than those for higher numbers of taxa when using at least 500 characters (Supplementary Figs. 3–6). Increasing the number of states per character has a major effect in improving accuracy and precision metrics for trees with few taxa, but a minor—albeit always positive—effect for trees with a high number of taxa. Considering the mean of metrics across simulation replicates, threshold values are passed at 500–5,000 characters for 5 taxa, at 100–500 characters for 10–50 taxa, and at 500 characters for 100 taxa or more (Fig. 2).

At tree length = 10, tree estimates for 20 or fewer taxa tend to be less accurate and precise than tree estimates for 50 or more taxa, keeping all other parameters equal (Fig.1, Supplementary Figs). For 5–10 taxa, not even 5,000 characters are sufficient to pass threshold values of the accuracy and precision metrics (except for the 4-states scenario, where 5,000 characters are sufficient for 10 taxa). Threshold values are passed at 500–5,000 characters for 20 taxa, at 100–500 characters for 50–100 taxa, and at 500 characters for 200 taxa (Fig. 2).

## 4. Discussion

Satisfactory levels of accuracy and precision of phylogenetic estimates require between 100 and 500 variable characters for most combinations of number of taxa, number of character states, and total tree length (Fig. 2). 50 characters are sufficient for 5–10 taxa with multistate characters evolving on shorter trees, while more than 500 characters are required for 200 taxa with shorter trees. It is worth stressing out that these numbers refer to an ideal scenario where the models used to infer a phylogeny are not misspecified, and they are relatively simple—phylogenetic analyses using more complex models likely require more characters. Additionally, simulated datasets had no missing data, which are typically common in empirical datasets—especially in datasets including extinct species—and can have a negative impact on phylogenetic estimates (32–35).

Empirical morphological datasets used for phylogenetic inference range in size between tens to hundreds (rarely up to few thousands) of characters. The mean number of characters observed in collections of several datasets usually varies between 35 and 150 (36–40), with datasets used for fossil tip-dating analyses tending to be larger (averaging 246 and 305 characters in two recent surveys (8,9)). Given that—under most scenarios considered here—more than 100 variable characters are needed to estimate phylogenies that are sufficiently accurate and precise, a substantial portion of empirical morphological datasets is likely too small to yield phylogenetic estimates that pass the thresholds of accuracy and precision set in this study. This result highlights the need for an increased research effort towards sampling more morphological characters for phylogenetic datasets. However, there are several limitations to the sampling of a large number of discrete morphological characters.

Building a morphological character matrix is an extremely time-consuming, laborious, and specialized task, whereby researchers must first identify morphological variability in observed phenotypes (trying to ignore variability that derives solely from ontogenetic, environmental, and preservational factors); then characterize that variability into definitions of each character and their possible states; then access relevant specimens—whether in-person or through digital metadata such as photographs and 3D models—to sample and score characters in the species within the studied clade. Additionally, there might be an intrinsic upper limit on the number of variable characters that can be characterized and scored in a given biological system. It is probably unrealistic to come up with hundreds of discrete morphological characters for structures such as gastropod shells (41), shark teeth (42), or palm fruits (43). The use of continuous characters in phylogenetics could represent a possible solution, as each continuous character can carry more information than each discrete character (for which the state space is limited). Moreover, several—if not most—discrete characters currently used in phylogenetics are arguably continuous characters that have been discretized, thus losing information about phylogenetically-informative features of the phenotype (44,45). While using continuous characters for phylogenetic inference comes with its own set of challenges (46,47), it represents a promising avenue for the future of morphological phylogenetics (48–51).

Regarding the impact of each variable considered in this study, the following general patterns emerge: adding characters improves phylogenetic estimates; more states per character slightly improve phylogenetic estimates, but this effect is mostly negligible on trees with more than 100 taxa; adding taxa worsens phylogenetic estimates on shorter trees, but can improve phylogenetic estimates on longer trees. A key finding of this simulation study is the major effect of tree length in the accuracy and precision of phylogenetic estimates. As tree length in unrooted trees is equivalent to the expected number of state changes per character across the tree, these results provide practical guidelines on which characters are better suited for phylogenetic analyses. A character that changes its state 10 times on a tree with 5 taxa is extremely uninformative due to multiple substitutions on the same branch erasing shared evolutionary history, while a similarly labile character on a tree with 200 taxa can support several clades across the phylogeny. These findings are coherent with previous literature on the beneficial effects of adding sampled taxa when substitution rates of characters are high, despite the added complexity in tree space (15–18). An intriguing consequence of these results is that, for phylogenies with more than 50 taxa, sampling homoplastic characters (characters that change multiple times independently) improves tree reconstruction compared to sampling strictly synapomorphic or autapomorphic characters (characters that change only once in the whole tree).

While this simulation study was designed to explore the effects of dataset size in morphological phylogenetics, the results shown here concern molecular phylogenetics as well. Molecular phylogenetic datasets tend to be much larger than morphological ones, usually ranging from thousands to millions of characters (or sites). However, when considering only variable characters, some kinds of molecular analyses—such as single gene tree reconstructions—might use datasets that are effectively much smaller in size, approaching the tens to few hundreds of characters typically seen in morphological studies (52,53). Depending on the number of variable sites, some genes might not carry enough information to reach the desired level of accuracy and precision for a phylogenetic estimate. This means that some discordance between gene trees—especially with a high number of taxa—might be at least partially attributed to stochasticity associated with data size and information content, rather than to biological processes such as incomplete lineage sorting and hybridization (54–56).

In conclusion, even under ideal conditions with no missing characters and no model misspecification, reasonably accurate and precise phylogenetic estimates require at least 100–500 variable characters, more than a substantial portion of empirical morphological datasets. These results outline the need for a dramatic increase in global research efforts (i.e., funding, hiring, academic training) geared towards the identification, characterization, collection, and digitization of interspecific morphological diversity (57–60), with the specific aim of building larger data matrices for phylogenetic studies. Without this added effort, downstream analyses that use morphological or total-evidence phylogenies to address macroevolutionary questions (e.g., diversification dynamics, phenotypic evolution) will often be based on faulty premises.

## Ethics

This work did not require ethical approval from a human subject or animal welfare committee.

## Data accessibility

Supplementary material is available online.

## Declaration of AI use

I have not used AI-assisted technologies in creating this article.

## Acknowledgements

I would like to thank Sebastian Höhna, Basanta Khakurel, and the rest of the Höhna Lab at LMU Munich for helpful discussions and feedback that improved this manuscript.

